# HGNChelper: identification and correction of invalid gene symbols for human and mouse

**DOI:** 10.1101/2020.09.16.300632

**Authors:** Sehyun Oh, Jasmine Abdelnabi, Ragheed Al-Dulaimi, Ayush Aggarwal, Marcel Ramos, Sean Davis, Markus Riester, Levi Waldron

## Abstract

Gene symbols are recognizable identifiers for gene names but are unstable and error-prone due to aliasing, manual entry, and unintentional conversion by spreadsheets to date format. Official gene symbol resources such as HUGO Gene Nomenclature Committee (HGNC) for human genes and the Mouse Genome Informatics project (MGI) for mouse genes provide authoritative sources of valid, aliased, and outdated symbols, but lack a programmatic interface and correction of symbols converted by spreadsheets. We present HGNChelper, an R package that identifies known aliases and outdated gene symbols based on the HGNC human and MGI mouse gene symbol databases, in addition to common mislabeling introduced by spreadsheets, and provides corrections where possible. HGNChelper identified invalid gene symbols in the most recent Molecular Signatures Database (mSigDB 7.0) and in platform annotation files of the Gene Expression Omnibus, with prevalence ranging from ∼3% in recent platforms to 30-40% in the earliest platforms from 2002-03. HGNChelper is installable from CRAN, with open development and issue tracking on GitHub and an associated pkgdown site https://waldronlab.io/HGNChelper/.

## Introduction

Gene symbols are widely used in biomedical research because they provide descriptive and memorable nomenclature for communication. However, gene symbols are constantly updated through the discoveries and re-identification of genes, resulting in new names or aliases. For example, *GCN5L2* (*G*eneral *C*ontrol of amino acid synthesis protein *5-L*ike *2*) is a gene symbol that was later discovered to function as a histone acetyltransferase and therefore renamed as *KAT2A* (*K*(lysine) *A*cetyl *T*ransferase *2A*)). [1] In addition to the rapid and constant updates on valid gene symbols, commonly used spreadsheet software, such as Microsoft Excel, mogrify some gene symbols, converting them into dates or floating-points numbers. [2,3] For example, ‘*DEC1*’, a symbol for ‘*D*eletion in *E*sophageal *C*ancer *1*’ gene, can be exported in date format, ‘1-DEC’. There have been attempts to rectify gene symbol issues, but they have largely been limited to Excel-mogrified gene symbols. Also the suggested solutions often reference static files with the corrections curated at the time of publication [3] or comprise scripts for detecting the existence of Excel-mogrified gene symbols without correction. [2] In recognition of the importance of the spreadsheet mogrification issues, HGNC recently announced that all symbols that auto-convert to dates in Excel have been changed. [4] However, much literature and public data still contains out-dated and incorrect gene symbols, motivating a convenient method of systematic detection and correction. To systematically identify historical aliases, correct for capitalization differences, and simultaneously correct spreadsheet-mogrified gene symbols, we built the HGNChelper R package. HGNChelper maps different aliases and spreadsheet-mogrified gene symbols to approved gene symbols maintained by The HUGO Gene Nomenclature Committee (HGNC) database. [5] HGNChelper also supports mouse gene symbol correction based on the Mouse Genome Informatics (MGI) database. [6]

## Materials and Methods

Human gene symbols are accessed from HGNC Database ftp site (ftp://ftp.ebi.ac.uk/pub/databases/genenames/new/tsv/hgnc_complete_set.txt) [7] and mouse gene symbols are acquired from MGI Database (http://www.informatics.jax.org/downloads/reports/MGI_EntrezGene.rpt). [6] Human gene symbol correction is processed in three steps. First, capitalization is fixed: all letters are converted to upper-case, except the open reading frame (orf) nomenclature, which is written in lower-case. Second, dates or floating-point numbers generated via Excel-mogrification are corrected using a custom index generated by importing all human gene symbols into Excel, exporting them in all available date formats, and collecting any gene symbols that are different from the originals. In the last and most commonly applied step, aliases are updated to approved gene symbols in the HGNC database. Mouse gene symbol correction follows the same three steps as in human gene symbol correction, except the capitalization step since mouse gene symbols begin with an uppercase character, followed by all lowercase.

## Results

The main function, checkGeneSymbols, returns a data frame with one row per input gene. The first column of the data frame shows the inputs, the second column indicates whether the symbols are valid, and a third column provides a corrected gene symbol where possible. The first line of the output indicates when the package’s built-in map was last updated. Because the gene symbol databases are updated as frequently as every day, we provide the getCurrentHumanMap and getCurrentMouseMap functions for updating the reference map without requiring an HGNChelper software update. These functions fetch the most up-to-date version of the map from HGNC and MGI, respectively, and users can provide the output of these functions through the map argument of checkGeneSymbols function. However, fetching a new map requires internet access and takes longer than using the package’s built-in index.

To evaluate the performance of HGNChelper, we quantified the extent of invalid gene symbols present in platform annotation files in the Gene Expression Omnibus (GEO) database from 2002 to 2020. We downloaded 20,716 GEO platform annotation (GPL) files using GEOquery::getGEO [8], of which 2,044 platforms were suspected to contain gene symbol information based on matching to valid symbols. There is a clear trend of increasing proportion of invalid gene symbols with age of platform submission (Fig 1), ranging from an average of ∼3% for recent platforms and increasing with age to ∼20% in 2010 and 30-40% in the earliest platforms from 2002-03. The overall proportion of the valid gene symbols was 79%, increasing to 92% after HGNChelper correction. The 8% of remaining, invalid gene symbols were mostly long non-coding RNA (lncRNA), pseudogenes, commercial product IDs such as probe ID, missing data, and gene symbols from non-human species, erroneously included together with human gene symbols. We also checked the validity of gene symbols in the Molecular Signatures Database (MSigDB 7.0). [9] Out of 38,040 gene symbols used in MSigDB version 7.0, 850 were invalid, and this number reduces to 453 after HGNChelper correction, of which the majority were lncRNA and a few withdrawn symbols.

**Fig 1.**
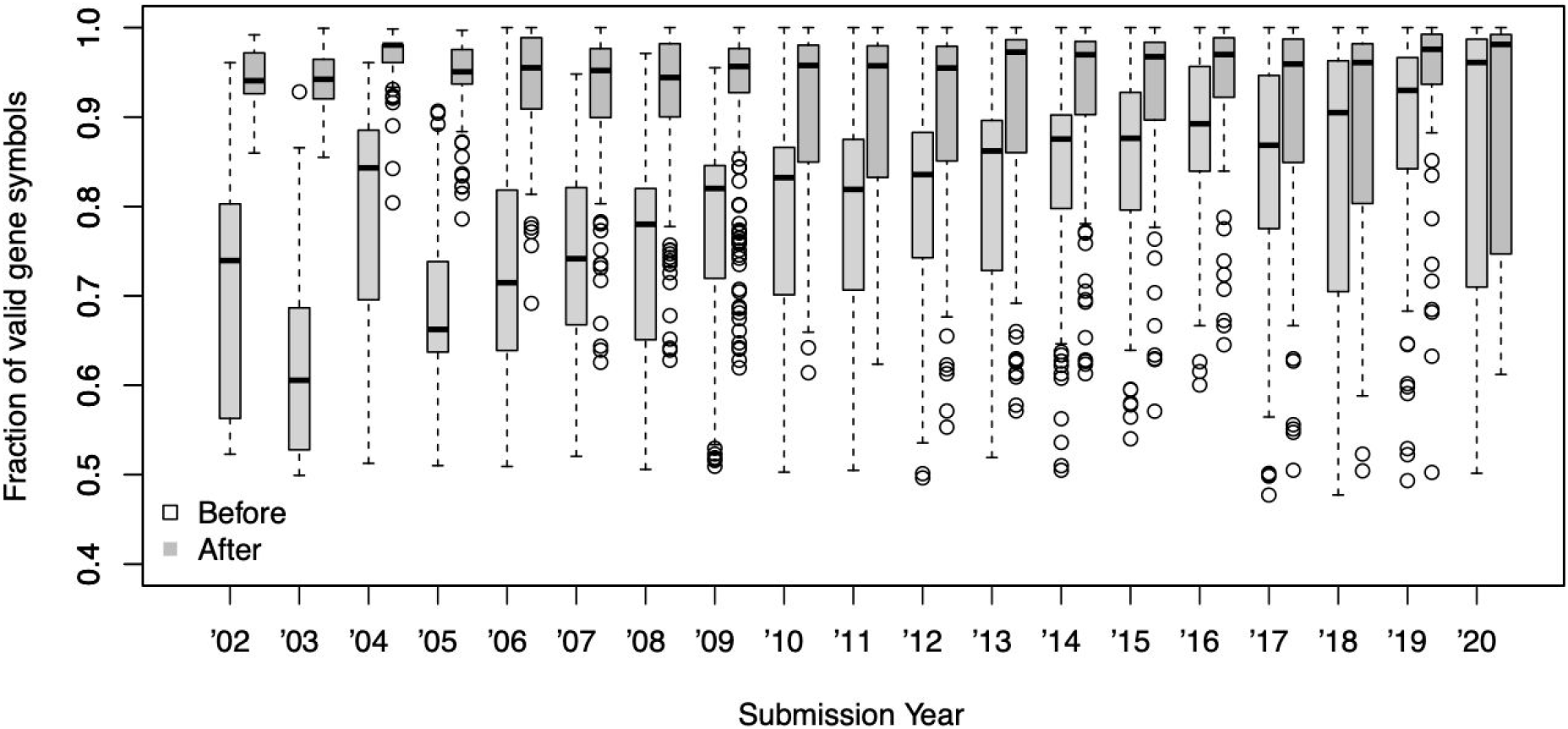
The fraction of valid gene symbols in GPL files grouped by year of data submission. Each dot represents a unique GPL. Older entries show a smaller fraction of valid gene symbols than more recent entries (Before, white box), but many of which are successfully corrected by HGNChelper (After, grey box).

## Discussion

Gene symbols are error-prone and unstable, but remain in common use for their memorability and interpretability. Our analysis of public databases containing gene symbols emphasizes the need for gene symbol correction particularly when using symbols from older datasets and reported results. Such correction should be routinely done when gene symbols are part of high-throughput analysis, such as re-analysis of targeted gene panels for precision medicine, which tend to be annotated with gene symbols (e.g. [10]), in Gene Set Enrichment Analysis using the gene symbol versions of popular databases such as MSigDB [9] or GeneSigDB [11], or when performing systematic review or meta-analysis of published multi-gene signatures (e.g. [12]). HGNChelper implements a programmatic and straightforward approach to the routine identification and correction of invalid gene symbols.

## Acknowledgements

This work was supported by the National Cancer Institute (NCI) grant U24-CA180996 to LW.

## References

1. Poux AN, Cebrat M, Kim CM, Cole PA, Marmorstein R. Structure of the GCN5 histone acetyltransferase bound to a bisubstrate inhibitor. Proc Natl Acad Sci U S A. 2002;99: 14065–14070.

2. Zeeberg BR, Riss J, Kane DW, Bussey KJ, Uchio E, Linehan WM, et al. Mistaken identifiers: gene name errors can be introduced inadvertently when using Excel in bioinformatics. BMC Bioinformatics. 2004;5: 80.

3. Ziemann M, Eren Y, El-Osta A. Gene name errors are widespread in the scientific literature. Genome Biol. 2016;17:p 177.

4. Bruford EA, Braschi B, Denny P, Jones TEM, Seal RL, Tweedie S. Guidelines for human gene nomenclature. Nat Genet. 2020;52: 754–758.

5. Yates B, Braschi B, Gray KA, Seal RL, Tweedie S, Bruford EA. Genenames.org: the HGNC and VGNC resources in 2017. Nucleic Acids Res. 2017;45: D619–D625.

6. Bult CJ, Blake JA, Smith CL, Kadin JA, Richardson JE, Mouse Genome Database Group. Mouse Genome Database (MGD) 2019. Nucleic Acids Res. 2019;47: D801–D806.

7. Home | HUGO Gene Nomenclature Committee. [cited 2 May 2020]. Available: http://www.genenames.org

8. Davis S, Meltzer PS. GEOquery: a bridge between the Gene Expression Omnibus (GEO) and BioConductor. Bioinformatics. 2007;23: 1846–1847.

9. Liberzon A, Subramanian A, Pinchback R, Thorvaldsdóttir H, Tamayo P, Mesirov JP. Molecular signatures database (MSigDB) 3.0. Bioinformatics. 2011;27: 1739–1740.

10. McCabe MJ, Gauthier M-EA, Chan C-L, Thompson TJ, De Sousa SMC, Puttick C, et al. Development and validation of a targeted gene sequencing panel for application to disparate cancers. Sci Rep. 2019;9: 17052.

11. Culhane AC, Schwarzl T, Sultana R, Picard KC, Picard SC, Lu TH, et al. GeneSigDB--a curated database of gene expression signatures. Nucleic Acids Res. 2010;38: D716–25.

12. Waldron L, Haibe-Kains B, Culhane AC, Riester M, Ding J, Wang XV, et al. Comparative meta-analysis of prognostic gene signatures for late-stage ovarian cancer. J Natl Cancer Inst. 2014;106. doi: 10.1093/jnci/dju049

